# Resting state periodic and aperiodic brain oscillations from birth to preschool years: Aperiodic maturity predicts developmental course

**DOI:** 10.1101/2025.05.11.653151

**Authors:** Heather L. Green, J. Christopher Edgar, Kylie Mol, Marybeth McNamee, Laura Prosser, Mina Kim, Emily S. Kuschner, Gregory A. Miller, Yuhan Chen

**Author notes:** Address for Correspondence: Yuhan Chen, Ph.D., The Children’s Hospital of Philadelphia 3401 Civic Center Blvd, Seashore House 1F, Room 116B Philadelphia, PA 19104. Shared first authors (Heather L. Green, J. Christopher Edgar).

## Abstract

The non-invasive assessment of resting-state (RS) neural activity via electrophysiology provides information on brain function and brain health. Our understanding of RS neural activity in older children and adults is limited by our poor understanding of the maturation of RS activity (oscillatory and non-oscillatory) from infancy to preschool ages. The present study used MEG with source imaging and an adapted Dark Room eyes-open task to assess oscillatory and non-oscillatory RS activity. 107 typically developing children 2 to 68 months were enrolled. For the Dark-Room eyes-open task, each child alternated between viewing an Inscapes video without audio for 20s and then resting with their eyes open for 30s in total darkness. This was repeated for 6 cycles. Whole-brain RS activity maps were computed using Minimum Norm Estimates, with RS power spectra divided into estimates of periodic measures (dominant frequency and power) and aperiodic measures (the exponent - slope of the 1/f function; the offset - vertical displacement of the 1/f function). An infant dominant peak was observed more often in the Dark-Room (94% of children) than in the Video-On condition (83% of children). The Dark-Room condition elicited a 36% increase in dominant oscillation activity than the Video-On condition. The maturation of the parietal-occipital periodic dominant frequency increased non-linearly as a function of age. The maturation of aperiodic measures decreased nonlinearly as a function of age, with aperiodic measures as well as their maturation rate differing across the brain. Finally, more mature aperiodic values predicted better adaptive behaviors and daily living skills. Present findings demonstrate that (1) the use of an appropriate Dark-Room eyes-open task provides measures of young child RS periodic activity with excellent SNR, (2) an understanding of the development of infant RS activity is best achieved via obtaining measures in brain source space in order to detect regional differences in aperiodic activity, and (3) a more mature aperiodic value predicts higher developmental behavior scores.

## Background

Brain development during embryonic and early fetal periods primarily involves the migration of neurons to specific locations, followed by events that establish a regional identity for each neuron [1].

Brain development during the late fetal period through infancy involves specification and refinement of local neural circuits and the formation of local and long-range connections between neurons [2]. Research on infant brain maturation is of interest given that the first months after birth are a peak period of neural reorganization, with neural maturation during this period associated with normal variation in developmental milestones as well as developmental delays, future mental illness, and cognitive impairment.

Using magnetoencephalography (MEG) combined with structural magnetic resonance imaging (MRI) and distributed source modeling to directly assess neural generator activity, infant and toddler studies have described the maturation of left and right auditory cortex encoding processes [3], documented the maturation of left and right primary visual and fusiform gyri activity when viewing faces [4], and shown that white-matter diffusion maturation near auditory cortex (supporting faster neural transmission), predicts decreasing latency of primary auditory cortex neural activity [5]. To interpret the maturation of primary sensory and higher-order neural processes as well as the behavior they support, it is critical to understand resting state (RS) neural activity, as infant RS neural activity is a foundation on which brain function and behavior emerge.

Literature dating to the 1940s demonstrates changes in RS neural activity from birth through adulthood, with an age-related decrease in delta and theta activity and an age-related increase in alpha, beta, and gamma activity [6–18]. Scientists examining the maturation of RS neural activity often study the dominant frequency, referred to in older children and adults as the ‘peak alpha frequency’ and in infants called the infant ‘alpha’ rhythm, or the ‘infant dominant frequency’ [19–23]. The dominant peak is observed at a lower frequency in infants and children than adults [6, 24–29], with dominant oscillation activity reflecting the excitability of a cortical region [30–33], associated with top-down cognitive control processes [34–36], and with an increase in the dominant frequency across development likely reflecting the ability to process information more quickly [13, 24]. Evidence supporting this latter association comes from studies showing that the dominant frequency predicts processing speed in neurotypical children [25, 37].

To date, challenges have hindered such research in infants and young children. In older child and adult electrophysiology RS studies, brain measures are often obtained in an eyes-closed condition, as the RS dominant peak is much stronger when the eyes are closed than open [13, 38–40]. Infants and young children however are unable to keep their eyes closed for an extended period on command and as a result, almost all infant/toddler RS studies acquire data while the child is at rest with the eyes open, and often while viewing visual stimuli [23, 41–46]. This strategy has limitations, as the dominant peak is much better identified in an eyes-closed than eyes-open condition [22, 42, 47–49]. For example, several EEG studies examining school-age children found a significantly larger dominant response in an eyes-closed than an eyes-open task [47–49]. A meta-analysis by Freschl et al. [50] highlights the difficulty in obtaining RS data in infants. A review of their Table 1 shows that of the included 3,882 subjects, only ∼5% were under 1 year old (with half of these from a single 1990 study), with most of the included infant studies reporting on eyes-open data.

**Table 1.** (A) Mean ± Standard Deviation of VABS domain standard scores. (B) Mixed-effects model main effects.

|  | ABC | Communication | Daily Living Skills | Socialization | Motor |
| --- | --- | --- | --- | --- | --- |
| **A. VABS domain scores** | 97.28 ± 8.51 | 99.43 ± 7.93 | 97.23 ± 10.36 | 98.70 ± 8.33 | 98.91 ± 10.06 |
| **B. Mixed model results** | ABC | Communication | Daily Living Skills | Socialization | Motor |
| **Aperiodic Exponent** | $F(1,732) = 52.46,$ | $F(1,732) = 2.87,$<br>$p = .27$ | $F(1,732) = 77.34,$ | $F(1,732) = 6.30,$<br>$p = .01$ | $F(1,732) = 8.86,$<br>$p = .004^*$ |
| | $p = .0001^{**}$ | | $p < .0001^{**}$ | | |
| **Aperiodic Offset** | $F(1,732) = 28.34,$<br>$p < .0001^{**}$ | $F(1,732) = 2.64,$<br>$p = .10$ | $F(1,732) = 68.22,$<br>$p < .0001^{**}$ | $F(1,732) = 0.89,$<br>$p = .35$ | $F(1,732) = 0.28,$<br>$p = .60$ |

Studies from the 1940s to the 1990s suggest that obtaining RS alpha measures in a dark room with eyes open is a viable alternative to the traditional RS eyes-closed exam [e.g., 23, 39, 51-55]. To explore this claim, a recent study from Edgar et al. [40] compared RS alpha obtained in eyes-closed, eyes- open, and eyes-open dark-room (DR) conditions (Dark-Room task described below in Methods) in neurotypical children and in children with autism spectrum disorder 6.9 to 12.6 years old. High consistency was observed for the eyes-closed and eyes-open Dark-Room alpha frequency and power measures (ICCs≥0.80) versus RS alpha activity near noise levels in the eyes-open condition. The present study assesses infant RS activity via the use of our Dark-Room RS task optimal for infants and young children.

Separate from that issue of data acquisition, a different methodological issue arises in data analysis and interpretation. Scientists have recently called into question previous electrophysiology RS findings, noting that most RS power-spectrum analyses conflate two brain processes. RS neural activity exhibits aperiodic background activity (contributing power across all frequencies when subjected to standard power-spectrum analysis) that co-exists with periodic oscillations [26, 27, 56]. In particular, RS electrophysiology power spectra show a characteristic 1/f power distribution (higher power at lower frequencies). Accordingly, RS analyses need to distinguish the 1/f <u>aperiodic</u> activity (often referred to as non-sinusoidal ‘background noise’ [57, 58] but very likely meaningful [56, 59, 60]) from periodic oscillatory activity to understand RS neural activity (for examples, see [56, 61]). A growing literature relies on the ‘specparam’ toolbox [56, 60] to separately parameterize periodic and aperiodic activity [59, 60, 62]. From infancy to late adolescence the RS 1/f power-spectrum slope flattens as a function of age until eventually stabilizing at an adult level [29, 42, 43, 62–65].

Unfortunately, most parameterization efforts to assess RS activity have been problematic, because most studies have conducted the analyses on EEG sensor data, thus assessing power spectra measures that reflect activity from multiple brain regions (for discussions of the limitations associated with this approach see [66–70]). The blurring of activity from multiple brain sources has often been even more severe, because in most studies aperiodic measures have been obtained from power spectra obtained via an average of EEG channels [43, 71] or via two or more regional clusters of EEG sensors [22, 41, 42, 44, 48, 71, 72]. These EEG cluster studies have produced inconsistent results.

In contrast to sensor space, EEG and MEG analysis in source space is designed to identify and distinguish regional sources. To our knowledge, only two pediatric studies have explored regional differences of the aperiodic exponent and offset measures in source space. Vandewouw et al. [45] undertook MEG source analysis to assess regional differences in age and aperiodic exponent and offset associations in 69 children and adults 1 to 38 years old. They found that across brain regions the aperiodic exponent decreased with age. Edgar et al. (in review) compared Dark-Room and Eyes-Closed MEG aperiodic measures in children 6.0 to 15.9 years old at 15 brain regions and showed high consistency between the Dark-Room and Eyes-Closed aperiodic exponent and offset parameter values across the brain regions. Edgar et al. also observed regional differences in the aperiodic values as well as regional differences in their maturation. With respect to brain development, a limitation of the above two studies is that Edgar et al. did not include young children, and in the Vandewouw et al. study there were no children under 1 year old and few children 1 to 3 years old.

Given regional differences in infant maturation of brain structure and chemistry, regional differences in RS activity are almost certain in very young brains. The present study sought to identify the beginnings of such RS differences, here examining infants and young children (2 months to 5 years old) and assessing RS activity in a relatively large sample (N=107). The present study addressed the possibility of regional differences in RS activity, as well as age-specific associations between RS activity and behavior.

Hypotheses were: (1) that in infants and young children Dark-Room data would provide more evaluable data and a better SNR than traditional video-stimulus data, (2) that source-space analyses would reveal regional differences in periodic and perhaps aperiodic activity as well as their maturation, and (3) that such aperiodic measures would predict behavioral development.

## Methods

### Participants

All infants and children were typically developing. Inclusion criteria included (1) no premature birth (<37 weeks gestation); (2) no concerns regarding developmental delay; (3) no history of seizure disorder; and (4) no known hearing or visual impairment. The study was approved by the Children’s Hospital of Philadelphia Institutional Review Board (IRB), and all families gave written consent.

Of 107 enrolled children, evaluable MEG data was acquired from 102 children (infants<36 months N=78, 31 females; young children 36 to 68 months N=24, 10 females). Five infants were excluded due to excess movement artifact during imaging (N=4) or technical difficulties (head position indicator (HPI) coil detached during exam N=1). Thirty-three of the 102 infants (13 females) had evaluable MEG data from more than one timepoint (2 Timepoints N=19; 3 Timepoints N=10; 4 Timepoints N=3; 6 Timepoints N=1). For the 33 infants with longitudinal data, only a single timepoint was included in the cross-sectional analyses, with in-house code used to randomly select one timepoint for each infant.

### RS Dark-Room Paradigm

For the RS Dark-Room eyes-open MEG task, over a 5-min period the child alternated between viewing an Inscapes video without audio for 20s [73] and then resting with their eyes open for 30s in total darkness, alternating 6 times. During the task, a parent and research assistant stayed in the room with the child. Figure 1A illustrates the Dark-Room paradigm.

**Figure 1.**
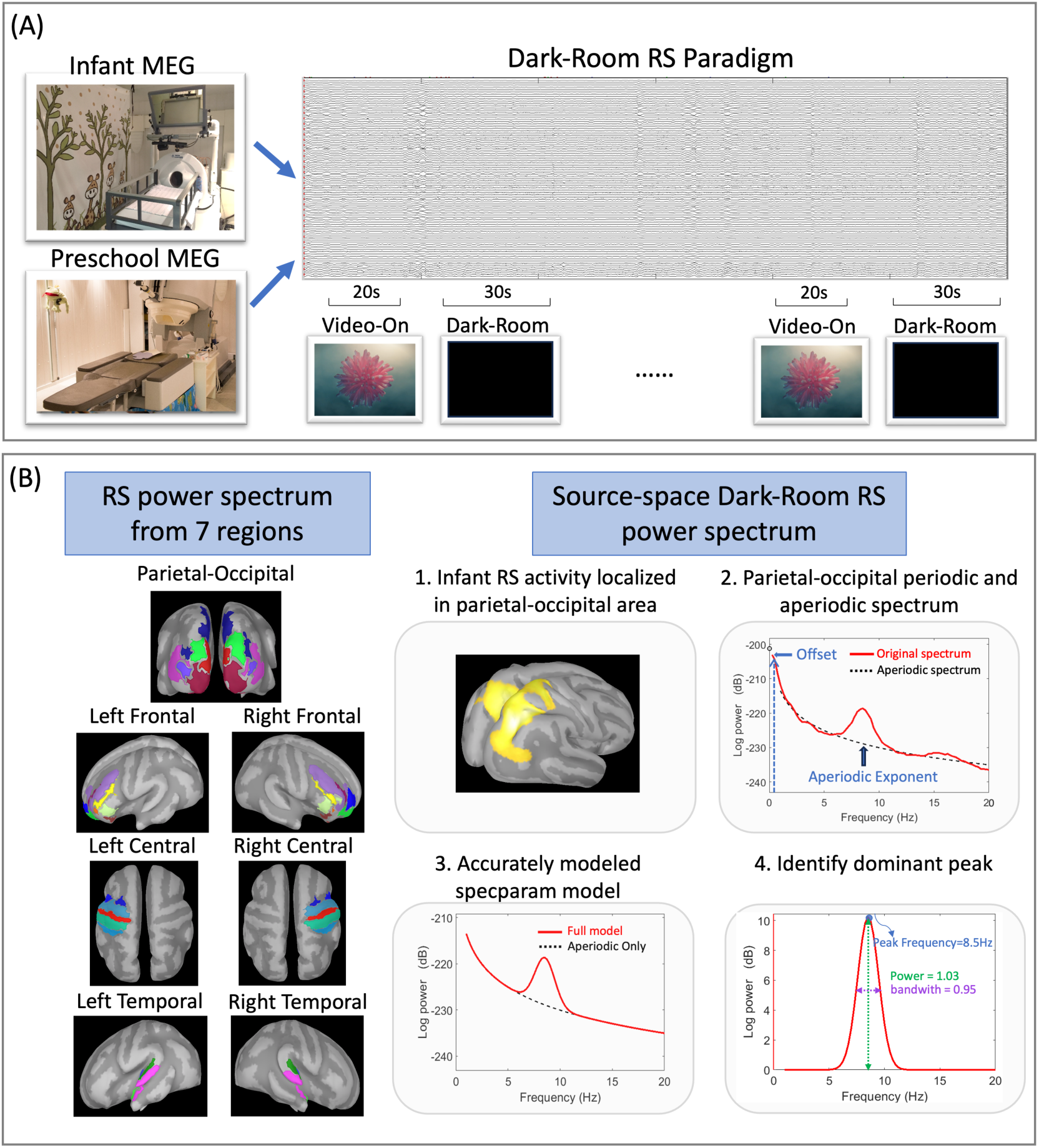
(A) Dark-Room RS task. Infant MEG data (children<36 months) were recorded using the 123- channel Artemis system. Preschool MEG data (children>36 months) were recorded using the 275-channel CTF system. (B) Source-space RS measures were obtained from seven regions of interest (ROIs; left side of panel B), with periodic and aperiodic measures obtained using the specparam algorithm. The right side of panel B demonstrates how specparam models features of the power spectrum via performing a sequential decomposition into periodic and aperiodic components. The periodic RS dominant rhythm parameters (center frequency, amplitude, and bandwidth) were obtained from a midline parietal-occipital region. RS aperiodic parameters (aperiodic exponent and offset) were obtained from all seven ROIs.

### MEG Data Acquisition

For children under 36 months, MEG data were recorded in a magnetically shielded room using an Artemis123 system (Tristan Technologies, Inc., San Diego, CA, USA) with a sampling rate of 5,000Hz and a 0.1Hz high-pass filter. Artemis123 was designed for use with children from birth to 3 years of age [74, 75]. The Artemis system has 123 first-order axial gradiometers and a helmet circumference of 50cm, which corresponds to the median head circumference of 36-month-old children in the United States.

Before MEG acquisition, a fabric cap with 4 HPI coils was placed on the child’s head. The child’s head shape, anatomical landmarks (nasion, right and left preauricular points), and locations of the HPI coils were digitized using a FastSCAN System (Polhemus, Colchester, VT). Head position was continuously monitored during MEG recording using the HPI coils.

MEG data from children 36 months and older were acquired in the same room using a 275- channel CTF system (VSM MedTech, Coquitlam, BC) with synthetic third-order gradiometer noise correction applied. CTF MEG data were acquired with a sampling rate of 1,200Hz. Before data acquisition, 3 HPI coils were placed on the child’s head. The child’s head shape, anatomical landmarks, and HPI coils were digitized, and head position was monitored using the HPI coils.

The Artemis123 and CTF MEG systems used the same hardware for stimulus presentation (projector, presentation computer). Children were scanned in the supine position, viewing the same visual stimuli at the same the viewing distance and visual angle. Several strategies helped keep the children calm and engaged during the MEG exams [5, 76].

### MEG Source Analysis

MEG data were analyzed using Brainstorm [77]; http://neuroimage.usc.edu/brainstorm). Digitized FastSCAN surface points representing the shape of the infant’s head (*>*10,000 points) were used to co-register each child’s MEG data to an age-appropriate infant or young child MRI template [78, 79] using an affine transformation that accommodated global scale differences between the child’s anatomy and the atlas. The Destrieux cortical parcellations were used [80].

MEG data were downsampled to 1,000Hz and band-pass filtered 0.3 to 55Hz, with a 60Hz notch filter. Heartbeat and eyeblink artifacts were removed via independent component analyses. Other artifacts (e.g., movement, muscle artifact) were visually identified and marked as artifact. Times when the child was not attending to the stimuli were manually removed (e.g., falling asleep). Average head movement across time for each child (displacement of the HPI locations compared to the starting head location) was computed for use in statistical analyses.

Whole-brain RS activity maps were computed using Minimum Norm Estimates [MNE; 81, 82- 84], with minimum-norm imaging estimating the amplitude of brain sources constrained to the cortex and with the current dipoles oriented normal to the local cortical surface [85]. An MEG noise covariance matrix was obtained from an empty room recording obtained immediately prior to each child’s MEG scan. MNE solutions were computed with normalization as part of the inverse routine, based on the noise covariance. Forward solutions were computed using an overlapping spheres model. MNE source estimation was computed and mapped to an age-appropriate MRI template.

### Obtaining Source-level RS Periodic and Aperiodic Parameter Values

Virtual time courses at each vertex in brain source space were derived from the MNE solution (∼15,000 vertices). At each vertex, an average RS power spectrum density was computed via Welch estimation for power density, using Hanning windows of 4s and 50% overlap, computed separately for the Dark-Room and Video-On conditions. To compute regional periodic and aperiodic activity, and to limit the number of statistical analyses, power spectra were averaged across vertices within seven ROIs defined in the Destrieux cortical atlas for the Dark-Room and Video-On conditions [80]: parieto-occipital, left frontal, right frontal, left central, right central, left temporal, and right temporal (Figure 1B right panel).

Figure 1B shows how RS periodic and aperiodic measures were obtained using the specparam algorithm [formerly known as FOOOF; 56, 59, 60]. Specparam performs a series of decompositions of the power spectrum and divides the power spectrum into periodic and aperiodic components (Figure 1B), so that the original spectrum is fitted with 1/f periodic activity, and then the aperiodic signal is subtracted. Specparam then sequentially estimates and fits peaks in the residual periodic spectrum as Gaussians and subtracts the Gaussians from the power spectrum to refit the 1/f aperiodic function. The final specparam model provides estimates of: (1) periodic measures including spectral peaks, with center frequency and power computed for each identified peak, and (2) aperiodic measures including the exponent (slope of the 1/f function) and the offset (vertical displacement of the 1/f function, see Figure 2A).

**Figure 2.**
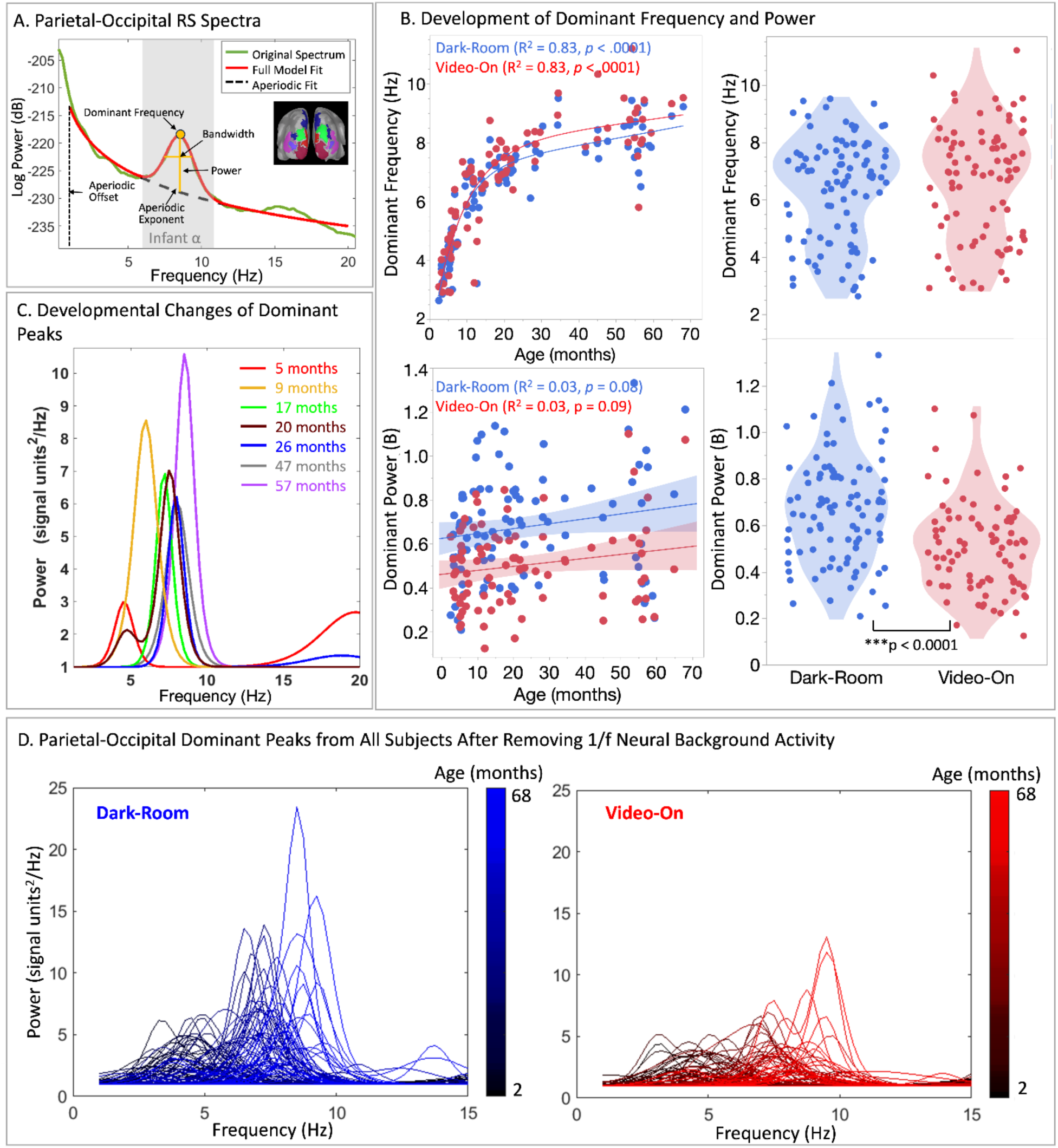
(A) Dark-Room power spectrum from a 3-year-old child and specparam fits. Specparam was used to identify the periodic dominant frequency and power for the midline parietal-occipital ROI and aperiodic exponent and offset for each of the seven ROIs. (B) Top left: The midline parietal-occipital dominant frequency increased exponentially as a function of age. Top right: Although Dark-Room and Video-Off conditions did not differ in parietal-occipital dominant frequency, the dominant frequency was observed more often in the Dark-Room (94%) than the Video-On condition (83%). Bottom left: Parietal- occipital dominant power was not associated with age. Bottom right: Dominant power was 36% stronger in the Dark-Room than the Video-On condition. (C) Examples of dominant peaks after removing 1/f aperiodic activity from seven representative children 5 months to 5 years old. (D) Midline parietal- occipital RS dominant rhythm spectrum for Dark-Room (blue) and Video-On (red) conditions for all subjects, after removing 1/f activity. The color indicates the age of each child (2 to 68 months) from deep blue/red (younger children) to light blue/red (older children).

Studies have shown that infant and young child dominant frequencies fall into a frequency range between 3 and 12Hz [22, 42, 72, 86–89]. The present study performed specparam models with the following settings: frequency range 1 to 20Hz, width for a detected peak between 1.5 and 8Hz, maximum number of detectable peaks 3, minimum peak height 3dB, a proximity threshold 2 standard deviations from the peak model, and fixed aperiodic mode. Model fit for each child was evaluated via the *R^2^* between the original spectra and fitted spectra for each condition and each ROI. The fitting procedure was very successful. Models with *R^2^*<0.80 were excluded (N=1).

Given that RS alpha activity is most prominent in parietal-occipital brain regions in older children and adults [90–92], RS dominant frequency and power were analyzed for the midline parietal-occipital ROI. Using in-house software, the dominant peak was identified as the largest peak in the parieto-occipital ROI between 2 and 12Hz. Supplementary Figure 1A shows several scenarios encountered when identifying the dominant peak. Specparam identified multiple peaks under 20Hz (maximum 3 peaks) in 49% and 47% of children in the Dark-Room and Video-On conditions, respectively (see Supplementary Figure 1B). For RS aperiodic analyses, and to evaluate whether there are regional differences in aperiodic activity, exponent and offset measures were recorded at all seven ROIs.

### Developmental Milestones

Developmental milestones were assessed via parent report using the Vineland Adaptive Behavior Scales [VABS-3; 93] for 53 children. Medical records were used to confirm typical development for 49 children whose caregivers did not complete the VABS-3. To evaluate associations between RS periodic and aperiodic neural activity and behavior, VABS-3 adaptive behavior composite (ABC) standard scores as well as four subdomain standard scores (*M*=100, *SD*=15) were used: communication, daily living skills, socialization, and motor.

### Statistical Analyses

Analyses were conducted using JMP Pro Version 18 (SAS Institute, Inc.). To evaluate associations between age and RS periodic activity (midline parietal-occipital dominant frequency and power), the Akaike information criterion (AIC) was used to determine whether a linear or non-linear model provided the better fit. To determine whether Dark-Room or Video-On provided better RS periodic signals, one-way ANOVAs tested the effect of condition on the periodic dominant frequency and power measures. A Fisher’s Exact Test evaluated whether a dominant peak was observed more often in the Dark-Room than the Video-On condition.

To assess changes in RS aperiodic activity (exponent, offset) as a function of age, AIC was used to determine whether a linear or non-linear model provided the better fit. To determine if there were regional differences in associations between age and aperiodic values, mixed-effect models were run separately for exponent and offset, with log age, ROI, head movement, condition, and log age X ROI interaction entered as fixed effects and subject entered as a random effect. Log age was used because of the nonlinear maturation of the aperiodic measures observed in the present sample (see Results). As age and MEG system (Artemis123, CTF) were highly correlated (younger children scanned using Artemis and older children scanned using CTF), MEG system was not included as a fixed effect in the mixed models.

Finally, relationships between RS measures and behavior were assessed. For RS periodic measures, each VABS-3 ABC and subdomain standard score entered as a dependent variable in separate analyses, with periodic dominant frequency (or power), condition, and periodic dominant frequency (or power) X condition interaction entered as independent variables. Age was entered as a random effect, allowing more nuanced analysis of age-related differences while accounting for the variability across subjects (i.e. allowing the age slope to vary across subjects). Mixed-effect models were run separately for dominant frequency and power, with a family-wise corrected threshold of *p*=.005 (*p*=.05 divided by 10 tests). That is a conservative threshold, because the 10 tests were not independent. For RS aperiodic measures, mixed-effect models were performed with each VABS-3 ABC and subdomain standard score entered as a dependent variable, and age entered as a random effect. Aperiodic measures (exponent or offset), ROI, condition, and their interactions were entered as independent variables, with family-wise correction again applied.

## Results

Across infant (Artemis123) and young child (CTF) samples, the mean (SD) amount of data collected for each condition was: Dark-Room (Artemis123) 125s (61), Dark-Room (CTF) 168s (32), Video-On (Artemis123) 88s (37), and Video-On (CTF) 112s (22). The total amount of artifact-free data was: Dark-Room (Artemis123) 102s (56), Dark-Room (CTF) 162s (32), Video-On (Artemis123) 73s (35), and Video-On (CTF) 105s (24).

As detailed in the Supplementary Figure 2 and Supplementary Table 1, to evaluate whether variables related to subject and data acquisition quality predicted specparam model fit (*R^2^*), a mixed-effect model was conducted with *R^2^* entered as the dependent variable, head movement, condition, age, ROI, and length of artifact-free data entered as fixed effects, and subject entered as a random effect. The specparam model fits were very good, with no difference in model fit between the conditions, and with model fit not associated with age or head movement. The only factor associated with specparam model fit was ROI (*p*<.0001), with midline parietal-occipital ROI having the best model fit (Supplementary Table 1). Supplementary Figure 3 shows an example of periodic RS activity at all seven ROIs in an 11-month-old child. Whereas in both conditions a large parietal-occipital dominant peak was quite evident, periodic peaks, if any, were much smaller and more difficult to identify in all other regions.

### RS Periodic Measures

As shown in Figure 2B, a dominant peak was observed more often in the Dark-Room (94% of children) than in the Video-On condition (83% of children; Fisher’s Exact Test *p*=.01). Comparison of a linear model (Dark-Room: AIC=300; Video-On: AIC=286) versus a nonlinear exponential growth model with 3 parameters (3P; Dark-Room: AIC=223; Video-On: AIC=234) showed that the nonlinear 3P model better represented the relationship between age and dominant frequency (Figure 2B). For the Dark-Room and Video-On conditions, the midline parietal-occipital dominant frequency rapidly increased during the first 2 years of life, with less rapid changes observed from 3 to 5 years old. The dominant frequency was ∼2.6Hz at 2 months and ∼9.5Hz at 68 months. Figure 2C illustrates the development of RS dominant peaks after removing the 1/f aperiodic activity from seven children 5 months to 5 years old.

The Dark-Room condition elicited higher dominant oscillation power (*M*=0.68 dB/Hz, *SD*=0.23) than the Video-On condition (*M*=0.50 dB/Hz, *SD*=0.19; *F*(1,179)=28.86, *p*<.0001, 95% CI (-0.11, -0.24); Figure 2B), with the Dark-Room condition providing a 36% increase in dominant oscillation activity. No association between age and dominant power was observed.

Figure 3 provides an example of how dominant peaks develop from a child scanned longitudinally from 3 to 39 months, with dominant frequency ranging from 3.97Hz at 3 months to 7.25Hz at 3 years.

**Figure 3.**
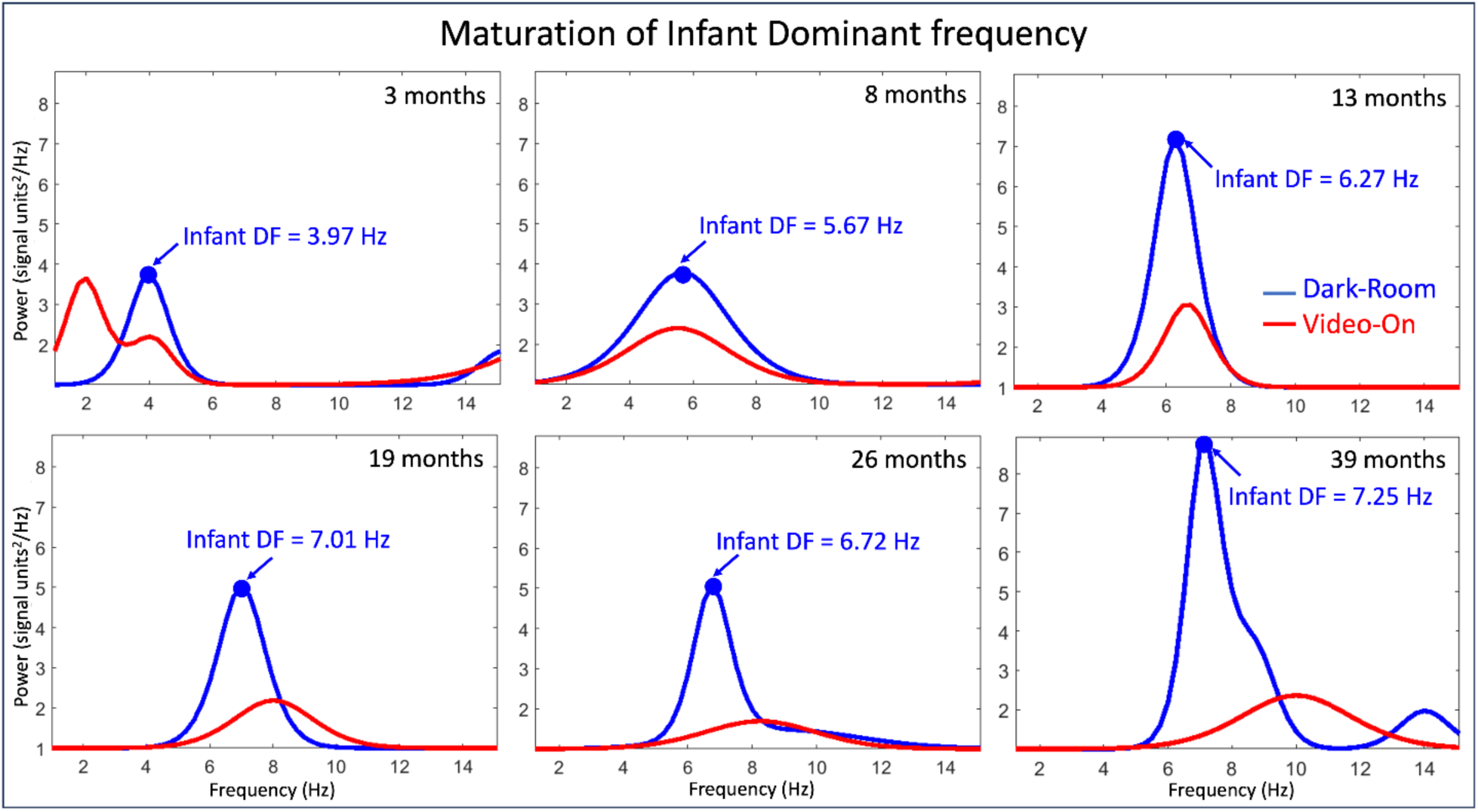
Maturation of dominant peak in one child. Dark-Room periodic activity is shown in blue and Video-On periodic activity in red. Blue arrows show the Dark-Room dominant frequency (DF).

### RS Aperiodic Parameter Values

Comparison of a linear model (Dark-Room: AIC=635; Video-On: AIC=563) versus a nonlinear 3P exponential growth model (Dark-Room: AIC=567; Video-On: AIC=547), showed that the nonlinear 3P model better represented the relationship between age and the aperiodic exponent. As shown in Figure 4A, rapid development of the Dark-Room aperiodic exponent was observed during the first year of life. A mixed-effects model showed main effects of log age (*F*(1, 102.3)=57.29, *p*<.0001) and ROI (*F*(1, 1213.2)=2.79, *p*=.01), as well as a log age X ROI interaction (*F*(1, 1212.4)=8.50, *p*<.0001). Simple-effect analyses of the ROI main effect showed that, compared to the midline parietal-occipital ROI, larger aperiodic exponent values were observed in the left central, left frontal, and left temporal regions (*ps*<.0001) and right central and right frontal regions (*ps*<.01).

**Figure 4.**
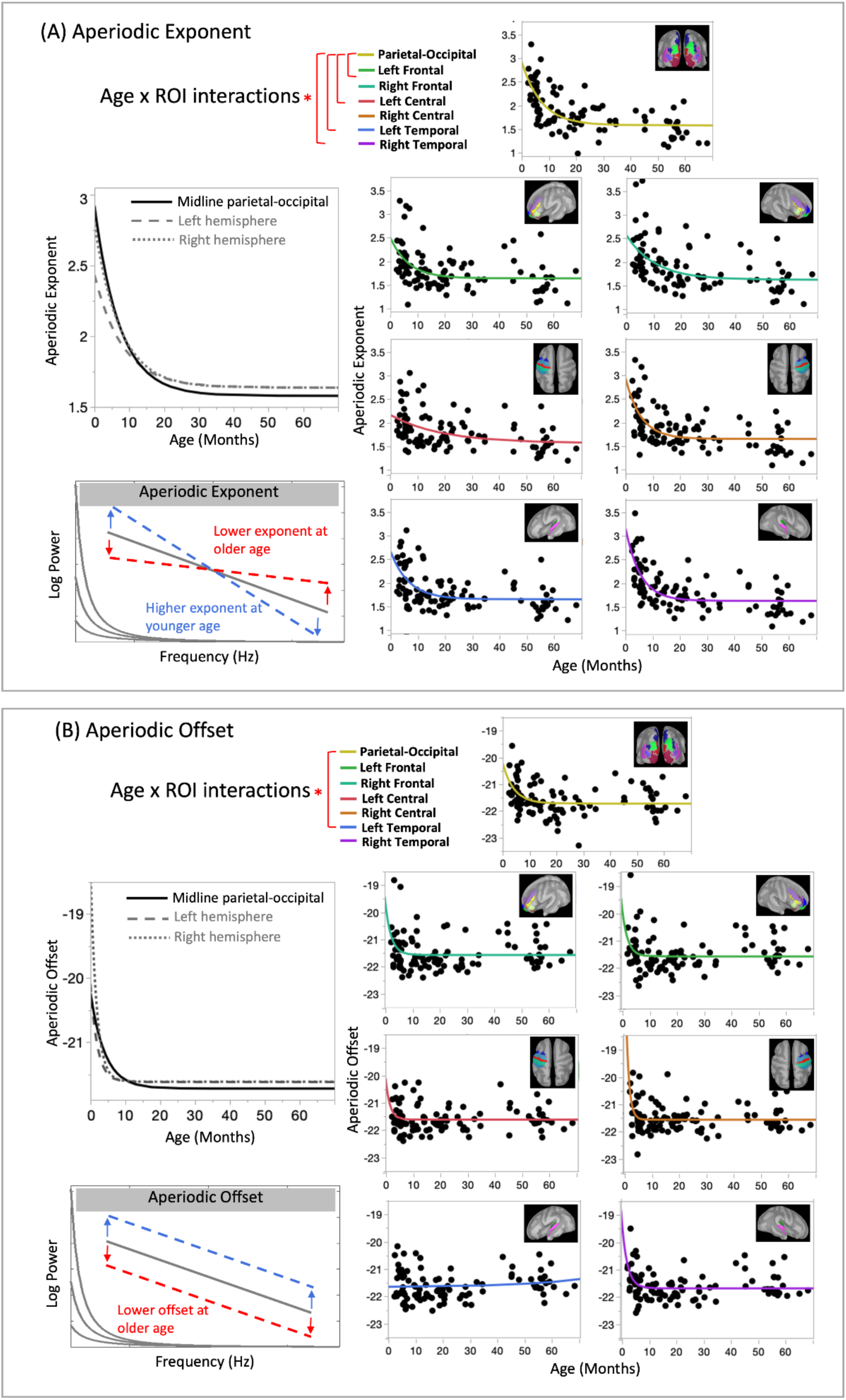
Given no main effect of condition, regionally specific developmental changes in aperiodic measures are shown only for the Dark-Room condition. **(A)** Top left: Aperiodic exponent by age. Bottom left: Illustration of the shift in the aperiodic exponent with age. Right: The relationship between the aperiodic exponent and age for each ROI. **(B)** Top left: Aperiodic offset by age. Bottom left: Illustration of the shift in the aperiodic offset with age. Right: The relationship between the aperiodic offset and age for each ROI.

Comparison of a linear model (Dark-Room: AIC=1174, Video-On: AIC=1059) versus a nonlinear 3P exponential growth model (Dark-Room: AIC=1150; Video-On: AIC=1050) showed that the nonlinear 3P model best represented the relationship between age and aperiodic offset. Figure 4B shows rapid aperiodic offset changes during the first months of life. Mixed-effect model analyses with aperiodic offset as the dependent variable showed effects of ROI (*F*(1, 1210.8)=3.76, *p*=.001) and a log age X ROI interaction (*F*(1,1210.0)=5.85, *p*<.0001). Simple-effect analyses of the ROI main effect showed that, compared to the midline parietal-occipital ROI, larger aperiodic offset values were observed in left frontal, left central, left temporal regions (*ps*<.0001) and in right frontal, right central, and right temporal regions (*ps*<.01). Head movement and condition did not account for significant variance in the aperiodic exponent or offset.

### Associations between Periodic and Aperiodic Parameter Values and Behavior

No association was observed between behavior ratings and dominant frequency or power in either condition (*p*s>0.05 after family-wise correction). Table 1A presents VABS-3 ABC and domain standard scores (N=53, 4 subjects excluded as multivariate outliers). RS aperiodic exponent and offset measures accounted for significant variance in domain-specific developmental scores. Table 1B shows that the aperiodic exponent accounted for significant variance in ABC, daily living skills, and motor domain scores. Aperiodic offset accounted for significant variance in ABC and daily living skills scores. For both aperiodic measures, smaller aperiodic values were associated with higher ABC and daily living skill scores. No associations were found between aperiodic measures and communication (language) and socialization scores. No interactions were observed between the fixed-effect variables. As an example, Figure 5 scatterplots show the associations between the aperiodic measures averaged across ROI and condition and VABS-3 daily living skill score, with *R^2^* and *p*-values based on zero-order correlations.

**Figure 5.**
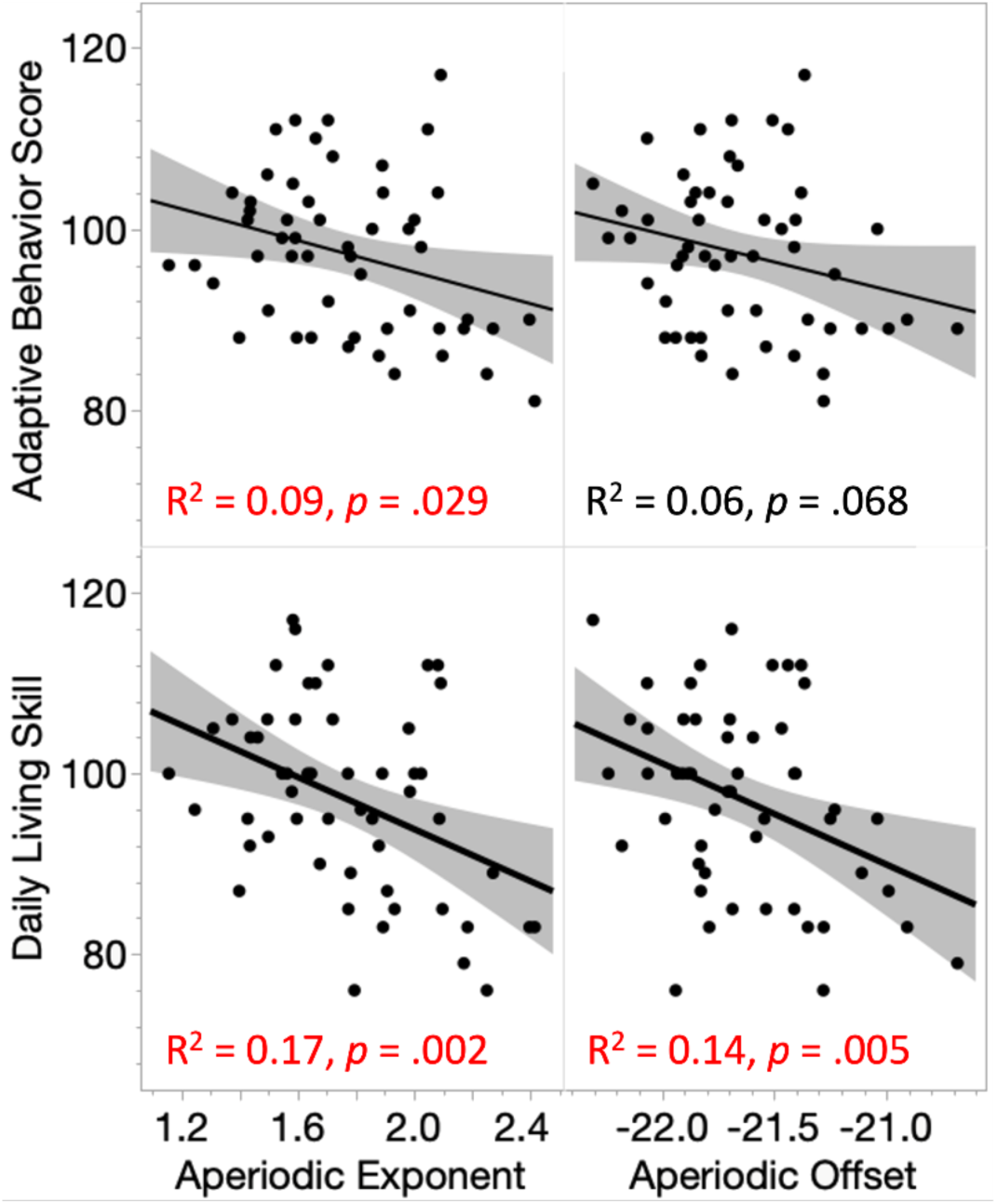
A more mature aperiodic measure predicts higher ABC and daily living skills scores.

## Discussion

Findings provide insight into the maturation of RS neural activity during the first years of life, replicating some recent reports while also pointing to region differences and behavior relationships not previously addressed. The expected age-related increase in the midline parietal-occipital dominant frequency was observed, with much more rapid maturation of the dominant rhythm in younger than older infants. Advantage for the Dark-Room versus Video-On condition was observed, with the dominant peak more often detected in the Dark-Room (94% of children) than the Video-On condition (83% of children), and with 36% higher dominant oscillation activity in the Dark-Room than Video-On condition. Aperiodic findings mirrored those from most previous studies, with exponent and offset values decreasing as a function of age. Findings support the small but growing literature in this area, and document both regional differences in aperiodic measures and regional differences in relationships between age and aperiodic measures. In addition, aperiodic measures were found to be associated with behavior ratings, with more mature aperiodic activity associated with higher adaptive behavior and daily living skills scores.

Findings thus demonstrate that (1) the use of an eyes-open Dark-Room task provides measures of young child RS periodic activity with an excellent SNR, (2) an understanding of the development of infant RS activity is best achieved via obtaining measures in brain source space in order to detect regional differences in aperiodic activity, and (3) a more mature aperiodic value predicts higher developmental scores.

### The Maturation of Midline Parietal-Occipital Periodic Neural Activity

As shown in Figure 2D, midline parietal-occipital dominant peaks were observed at all ages, with a dominant frequency observed at 2.6Hz at 2 months. The infant RS literature shows mixed results regarding the age when the dominant peak is first observed, as well as the pattern of age-related changes in dominant oscillation frequency and power. Factors likely accounting for mixed young child findings include: (1) examining RS periodic activity with [41–43, 46, 72] versus without having removed power contributed by 1/f aperiodic activity [23, 94], (2) examining RS activity in sensor space [23, 41–43, 46, 72, 95] versus source space [45], and (3) examining RS activity using an RS eyes-open task [41–43, 45, 46, 72, 95] versus a RS dark-room task [23]. As discussed below, present findings demonstrate advantages of both obtaining Dark-Room data and examining RS activity in source space.

Studies have reported an age-related increase in the dominant frequency in infants as young as 2 months [22, 23, 45-48, for meta-analysis see 50]. This pattern is often but not always observed. For example, Rico-Pico [72] observed a quadratic relationship between the dominant frequency and age, with an early increase, then a plateau, and then a decrease. Wilkinson et al. [46] reported an age-related increase in the dominant frequency in children only 6 months and older [46, see Figure 2 in Wilkinson et al.].

In the present study, a focus on midline parietal-occipital RS neural generators provided periodic dominant peaks that were easily detected in almost all infants, with a comparison of the Dark-Room and Video-On power spectrum used to identify the dominant peak when multiple periodic peaks were observed. As shown in Figure 2D and Supplementary Figure 1, a stronger periodic peak in the Dark- Room than Video-On condition allowed the periodic dominant peak to be easily identified, with the frequency of the dominant peak increasing as a function of age. As shown in Figure 3, this age effect was observed at the level of an individual infant. Present findings thus provide strong evidence for the maturation of the dominant oscillation from delta and theta ranges to the low alpha range during the first years old life. Present findings also demonstrate that the periodic dominant peak would often be difficult to detect in a Video-On condition, with the peak smaller and sometimes absent.

There is an ongoing conversation with respect to defining the infant and young child RS dominant oscillation. As reviewed in Stroganova et al. [23], some consider the delta- and theta-band activity observed in children a precursor of adult eyes-closed alpha activity [96–98]. Others, however, propose that RS alpha activity undergoes a developmental change different from delta and theta waves [17, 99]. As in Stroganova et al. [23], the present study adopted a “functional topography” approach [100], with the determination of a child dominant peak based on two main criteria: spatial distribution (larger in posterior than anterior brain regions) and functional reactivity (changes as a function of visual input, emerging given a homogeneous visual field [54]). Present findings support this functional topography approach. First, Supplementary Figure 3 demonstrates regional differences in the dominant peak, with largest dominant peaks observed in parietal-occipital regions and absent in frontal regions.

Such findings suggest that analysis strategies that average RS activity across EEG sensors risk diminishing or averaging out the dominant oscillation response. Second, as demonstrated in Supplementary Figure 1, in children with multiple periodic peaks, a comparison of Dark-Room and Eyes- Closed power spectra facilitated identification of the periodic peak as the peak functionally reactive to visual input (the greater synchronization of oscillatory activity given a homogenous visual field) and thus to confirm the Dark-Room periodic peak as the dominant peak. Differences between study findings are likely due to whether the RS dominant frequency and power identified via a functional topography approach used in our study, or via averaging periodic spectrum across all EEG sensors. The later approach could result in dominant peaks observed in fewer infants, as well as a shift of dominant frequency range (see Supplementary Figure 3).

With respect to the RS dominant power findings, there was only a trend finding of an age-related increase in midline parietal-occipital periodic dominant power (*p*=.08 for Dark-Room and *p*=.09 for Video-On condition). Dominant power findings are mixed, with some studies reporting an increase with age [23, 46, 71], and others reporting no association with age [22, 45, 48], these differences again are likely due to the differences noted above in how infant RS data are obtained and analyzed. Present findings suggest an age-related increase in dominant power, but a small effect size requiring a large sample for statistical confirmation.

In sum, present findings demonstrate that a RS Dark-Room paradigm and infant source-space RS analyses are well suited to assess the maturation of midline parietal-occipital RS periodic neural activity in infants and young children.

### Aperiodic RS Neural Activity

Present findings showed regional differences in the aperiodic measures as well as associations with age. Specifically, a nonlinear age-related decrease in the RS aperiodic measures was observed, with the aperiodic offset decreasing rapidly from birth and then plateauing at around 10 months, and with the aperiodic exponent decreasing rapidly from birth and then plateauing around 20 months.

Whereas most infant studies report age-related decreases for the aperiodic exponent [22, 42, 44, 45, 48, 71, 72, but see 46] and offset [22, 44, 45, 48, 62, 71, but see 46], neonate mice and human neonates show an increase in the 1/f exponent [101]. Chini et al. [101] hypothesized that the neonate-to-infant change in the direction of the 1/f exponent is explained by the decline of brain GABA levels [102] and cortical inhibition [103]. In children 1 to 7 months, it is also hypothesized that a wave of interneuronal death around this age [104] might induce an early shift from an E:I ratio decrease to an E:I ratio increase. This change might also reflect maturational changes in the integration of inhibitory interneurons into cortical circuity, with inhibitory circuits dominated by somatostatin-positive inhibitory interneurons during the first postnatal weeks [105, 106] and with parvalbumin-positive inhibitory interneurons later integrated [105–107].

After the first weeks of life, an age-related decrease in the aperiodic exponent and offset values (flattened RS power spectrum; Figure 4A & 4B bottom left plot) have been hypothesized to be associated with an age-related increase in the neural-circuit E:I ratio, with changes to the neural-circuit E:I due to less neural noise in older than younger children given an age-related increase in inhibitory GABAergic and an age-related decrease in glutamatergic signaling [108]. A relatively rapid decrease in the E:I ratio during the first years of life has been hypothesized to facilitate efficient information processing [48]. The present finding of a steeper RS power spectrum suggests more inhibition-dominant (I E) RS neuronal background in younger than older children. The above hypotheses are supported by findings from Edgar et al. (in preparation) that higher midline parietal-occipital GABA concentration (obtained using MRS) is associated with a less steep RS exponent.

In the present study, the association between age and the aperiodic measures differed across brain region, suggesting regionally-specific changes in the E:I balance from birth to 5 years. Such findings indicate the need to develop nuanced models of the maturation of infant neural activity. As an example, regional differences in the aperiodic measures are likely associated with regional differences in the maturation of gray and white matter, with some brain areas maturing relatively quickly [e.g., face processing 4, e.g., visual system 109], whereas others brain regions mature more slowly [e.g., auditory system 3, 5].

### RS Aperiodic Parameter Values Predict Behavior

In the present study, more mature aperiodic exponent and offset values were associated with higher adaptive behavior, daily living skills, and motor ability scores. Although aperiodic activity was observed to be regionally specific, the aperiodic and behavior associations were observed in all ROIs. This may reflect a general pattern, with a lower aperiodic exponent/offset likely reflecting less inhibitory and more excitatory activity (i.e., an increased E:I ratio) in the thalamocortical network to sensory input thus aiding in the development of better daily living skills and motor ability. The observed associations indicate moderate effects, with the aperiodic measures accounting for 9 to 17% of the variance in behavior scores.

In summary, present findings demonstrate that (1) the use of an appropriate Dark-Room eyes- open task provides measures of young child RS periodic activity with an excellent SNR, (2) an understanding of the development of infant RS activity is best achieved via obtaining measures in brain source space in order to detect regional differences in aperiodic activity, and (3) a more mature aperiodic value predicts higher developmental behavior scores.

## Acknowledgments

This work was supported by the National Institute of Child Health and Human Development to Dr. Edgar (R01HD093776) and Dr. Chen (3UG1HD068244-13S1, principal investigator Dr. Sara DeMauro); the National Institute of Mental Health to Dr. Edgar (R01MH10750, R21MH098204) and Dr. Chen (K01MH108822); the Nancy Lurie Marks Family Foundation to Dr. Green (principal investigator Roseann Schaff); and the Eagles Autism Foundation to Dr. Chen (Pilot Grant).

## Author Contributions

**HLG** – Formal analysis, Conceptualization, Data Curation, Investigation, Supervision, Writing- Original Draft, Review & Editing; **JCE** – Conceptualization, Statistical Analysis, Funding Acquisition, Supervision, Writing- Original Draft, Review & Editing; **KM** – Investigation; **MM** – Investigation; **LP** – Funding Acquisition; **MK** – Investigation, Writing - Review & Editing; **ESK** – Investigation; **GAM** – Writing - Review & Editing; **YC** – Conceptualization, Investigation, Formal Analysis, Methodology, Visualization, Supervision, Data Curation, Funding Acquisition, Writing – Original Draft & Editing. All authors read and approved the final manuscript.

## Data Availability Statement

The datasets generated and/or analyzed will be made available upon completion of this study and reasonable request.

