## Supplementary for "Resting state periodic and aperiodic brain oscillations from birth to preschool years: Aperiodic maturity predicts developmental course"

* Shared first authors

**Supplementary Materials include:**

Supplementary Figures 1 to 3.

Supplementary Table 1.

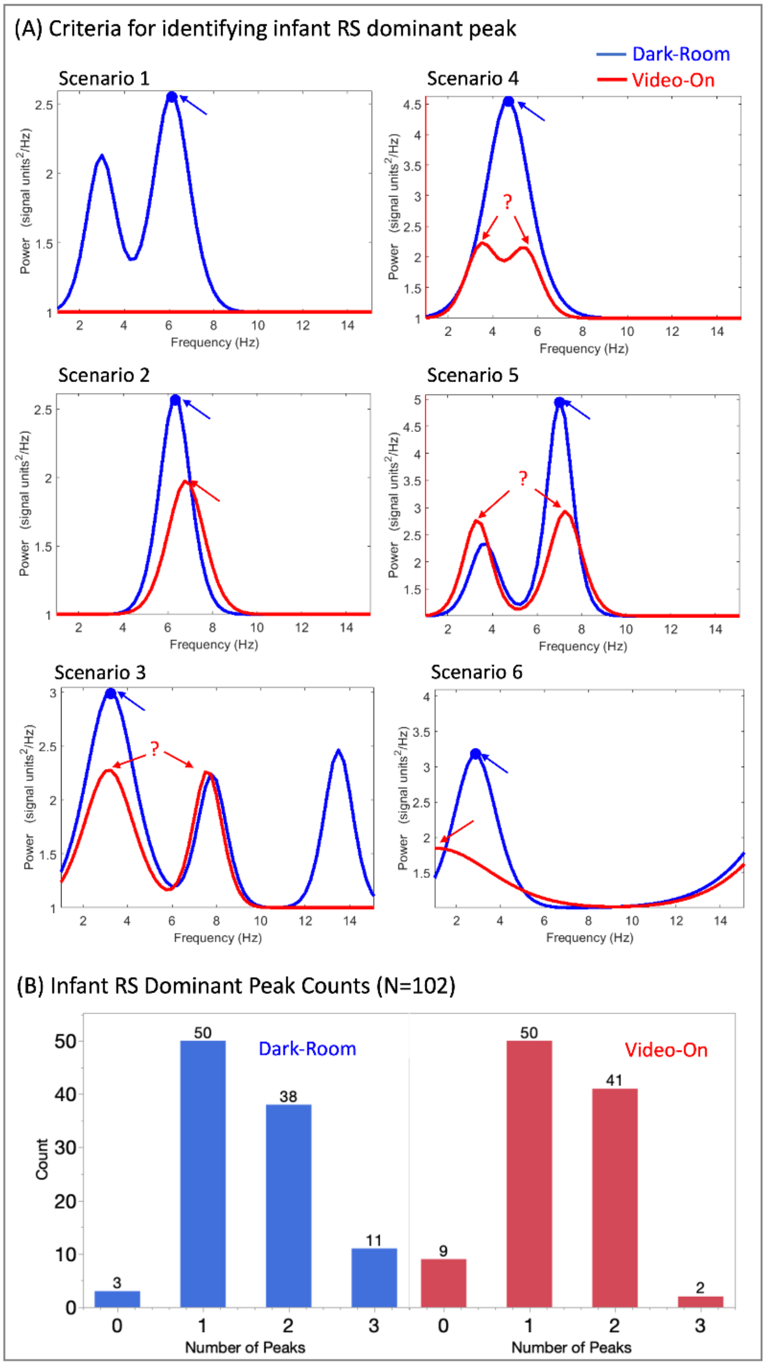


**Supplementary Figure 1**. (A) Identifying the periodic RS dominant peak. Blue arrows identify the Dark-Room dominant peak. Red arrows identify Video-On dominant peak. Question marks indicate cases where a Video-On dominant peak is difficult to identify due to similar amplitudes at different frequencies. For example, Scenario 1 shows that no periodic peak was observed in the Video-On condition, and 2 periodic peaks were observed in the Dark-Room condition, with the arrow pointing to the largest peak between 2 and 12Hz. Scenario 2 shows 1 periodic peak in both the Dark-Room and Video-On conditions. Scenarios 3, 4, and 5 show that, whereas similar amplitudes were observed for the 2 periodic peaks in the Video-On condition, only 1 large periodic peak was identified in the Dark-Room condition. Scenario 6 shows that the periodic peak identified in the Video-On condition was noise versus a clear periodic peak observed in the Dark-Room condition. The ages of representative children for each scenario are as follows: Scenario 1 (27 months), Scenario 2 (20 months), Scenario 3 (3 months), Scenario 4 (5 months), Scenario 5 (16 months), Scenario 6 (3 months). (B) RS periodic peak counts identified for each condition. For example, in the Dark-Room condition, from 1 to 20Hz, 2 peaks were identified in 38 children, and 3 peaks were identified in 11 children.

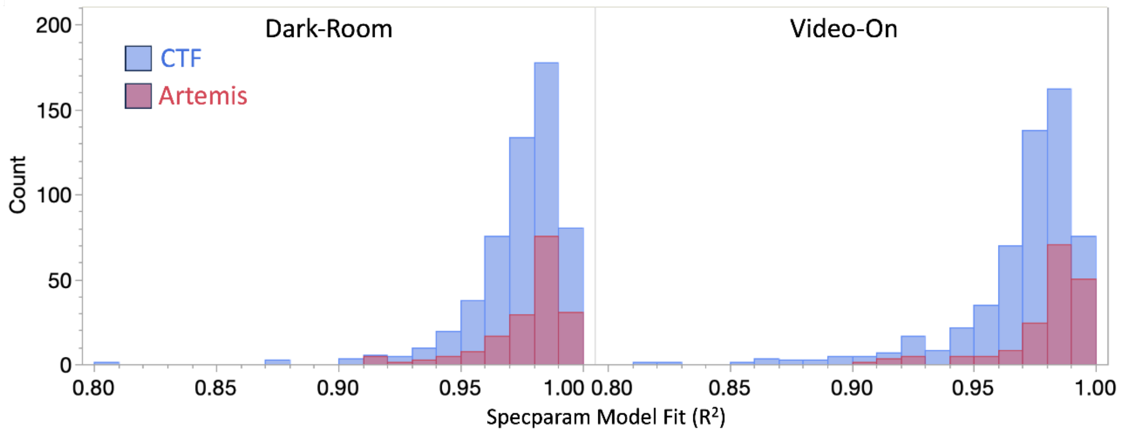


**Supplementary Figure 2.** Histograms of Specparam $R^2$ model fit (collapsing across ROIs) for the Dark-Room and Video-On conditions. Data from Artemis are shown as pink, and data from CTF are shown in blue. Only ROI accounted for variance in $R^2$ ($p$<.0001), with the midline parietal-occipital ROI having the best model fit ($M$ $R^2$=0.98, range 0.91 to 0.99). Simple-effect analyses showed significantly smaller $R^2$ values in left frontal, left central, and left right frontal ROIs than in the midline parietal-occipital ROI. Dark-Room and Video-On conditions did not differ in $R^2$ fit or proportion of artifact-free data (length of artifact-free data/length of total data collected per condition) ($ps$>.05). No association between age and head movement was observed ($p$=.25). Age was positively associated with proportion of artifact-free data for the Dark-Room and Video-On conditions ($ps$<.0001).

**Supplementary Table 1**. Descriptive statistics (mean (*M*), standard deviation (*SD*), minimum (*Min*) and maximum (*Max*) of specparam $R^2$ model fit for each ROI averaged across condition.

| **(B) Descriptive Statistics for Specparam Model Fit ($R^2$)** | | | | |
|---|---|---|---|---|
| **ROI** | ***M*** | ***SD*** | ***Min*** | ***Max*** |
| **Parietal-Occipital** | 0.98 | 0.02 | 0.91 | 0.99 |
| **Left Central** | 0.98 | 0.03 | 0.81 | 0.99 |
| **Right Central** | 0.98 | 0.02 | 0.90 | 0.99 |
| **Left Temporal** | 0.97 | 0.02 | 0.87 | 0.99 |
| **Right Temporal** | 0.98 | 0.02 | 0.91 | 0.99 |
| **Left Frontal** | 0.97 | 0.02 | 0.86 | 0.99 |
| **Right Frontal** | 0.97 | 0.02 | 0.81 | 0.99 |

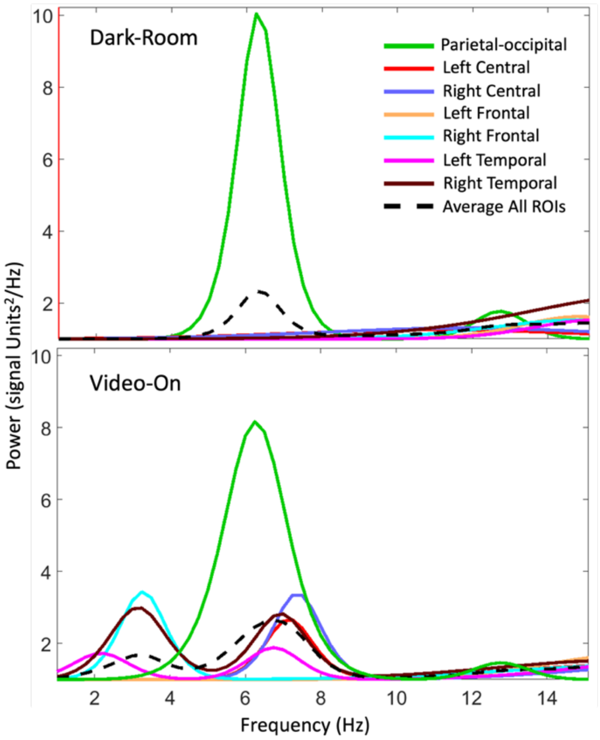


**Supplementary Figure 3**. Infant periodic peaks overlayed for all seven ROIs from a representative 11-month-old infant for the Dark-Room (top) and Video-On (bottom) conditions. The black dashed line shows periodic activity averaged across the seven ROIs. Whereas in both conditions a large parietal-occipital dominant peak was quite evident, periodic peaks, if any, were much smaller and more difficult to identify in all other regions. The use of a functional topography approach in the present study likely explains some of the differences between the results obtained here and other studies. As shown in this figure, two peaks were observed in the periodic spectrum averaged across ROIs (black dash line) in the Video-On condition. We hypothesized that this averaged response is similar to that obtained when averaging across EEG sensors. In particular, the observed average response is similar to that observed in Wilkinson et al. [46] and perhaps explains why in their study two periodic peaks were observed in some children: 69% of the infants 2 to 4 months old had a peak in the theta range (4–6Hz) and the alpha range (6.5–12Hz). As discussed in Stroganova et al. [23], it is likely that in infants a lower-frequency peak reflects the child's dominant oscillation activity generated by parieto-occipital neural generators, whereas a higher frequency at central EEG sensors reflects central rhythmic activity (in adults considered the adult sensorimotor mu rhythm). To this point, also of note are the Groppe et al. [110] ECoG findings, with substantial variability in the dominant frequency observed across cortical regions, and with theta-band activity (4-8Hz) the dominant oscillation in frontal, central, and temporal regions versus alpha-band activity largely limited to midline parietal and occipital regions. Although a thorough examination of regional differences in RS periodic activity was beyond the scope of the present study, the exploratory findings in Supplementary Figure 3 indicate the value of regionally specific measures of RS periodic activity.
